# A Distributed Computing Solution for Privacy-Preserving Genome-Wide Association Studies

**DOI:** 10.1101/2024.01.15.575678

**Authors:** Cláudia Brito, Pedro Ferreira, João Paulo

## Abstract

Breakthroughs in sequencing technologies led to an exponential growth of genomic data, providing unprecedented biological insights and new therapeutic applications. However, analyzing such large amounts of sensitive data raises key concerns regarding data privacy, specifically when the information is outsourced to third-party infrastructures for data storage and processing (*e*.*g*., cloud computing). Current solutions for data privacy protection resort to centralized designs or cryptographic primitives that impose considerable computational overheads, limiting their applicability to large-scale genomic analysis.

We introduce Gyosa, a secure and privacy-preserving distributed genomic analysis solution. Unlike in previous work, Gyosa follows a distributed processing design that enables handling larger amounts of genomic data in a scalable and efficient fashion. Further, by leveraging trusted execution environments (TEEs), namely Intel SGX, Gyosa allows users to confidentially delegate their GWAS analysis to untrusted third-party infrastructures. To overcome the memory limitations of SGX, we implement a computation partitioning scheme within Gyosa. This scheme reduces the number of operations done inside the TEEs while safeguarding the users’ genomic data privacy. By integrating this security scheme in *Glow*, Gyosa provides a secure and distributed environment that facilitates diverse GWAS studies. The experimental evaluation validates the applicability and scalability of Gyosa, reinforcing its ability to provide enhanced security guarantees. Further, the results show that, by distributing GWASes computations, one can achieve a practical and usable privacy-preserving solution.

## 1 Introduction

With the advent of next-generation sequencing (NGS) technologies, the cost of genome sequencing has decreased significantly, enabling the generation of large amounts of genomic data in a timely and cost-effective manner [33]. The availability of large-scale datasets opens up new avenues for research on genetic factors, but it also requires computationally efficient algorithms capable of handling the sheer magnitude of data.

Genomic Wide-Association Studies (GWAS) test the association of hundreds of thousands to millions of genetic variants in a cohort of individuals and find the variants that are statistically associated with a specific trait or disease [3, 36]. By 2021, more than 5700 GWASes have been conducted using data from more than one million individuals for more than 3300 traits [36].

However, the feasibility of running these algorithms and statistical methods relies on the existing computational power. When working on a single workstation, commonly called a server, users face limitations imposed by the existing computational capacity. The computational demands of GWAS are directly linked to variables like the number of genetic variants, the number of individuals, and the tested traits and phenotypes. For very large datasets, a single server’s computing and storage power may be insufficient. The conventional approach of enhancing server capacity by augmenting core processors, memory, and storage resources may encounter significant challenges, including exponentially escalating costs and inherent limitations associated with the presently available hardware [31]. Distributed computing, which allows using several servers (clusters of servers) in parallel, is a viable solution to reduce the runtime execution of parallelizable and data-intensive algorithms, such as GWAS [7]. Over the years, this paradigm has improved the parallelization and distribution of large-scale computations and their access to large amounts of data. However, acquiring, maintaining, and managing a distributed server infrastructure is a costly and complex task requiring high-end hardware and specialized human resources [31]. An accessible option involves running GWAS analysis remotely on distributed infrastructures managed by third-party entities, such as those provided by Cloud Computing (*e*.*g*., GCP,) and HPC services (*e*.*g*., TACC [35]). This approach may offer significant benefits, especially in scenarios where: *i*) a single entity, such as a Hospital, possesses a large genomic dataset but lacks the processing and storage power for analysis; and *ii)* a consortium of entities, including Hospitals and research laboratories (see Figure 1 in Supplementary Material), aims to collectively analyze their datasets in a unified remote infrastructure, enabling large-scale analysis using shared data.

**Fig. 1:**
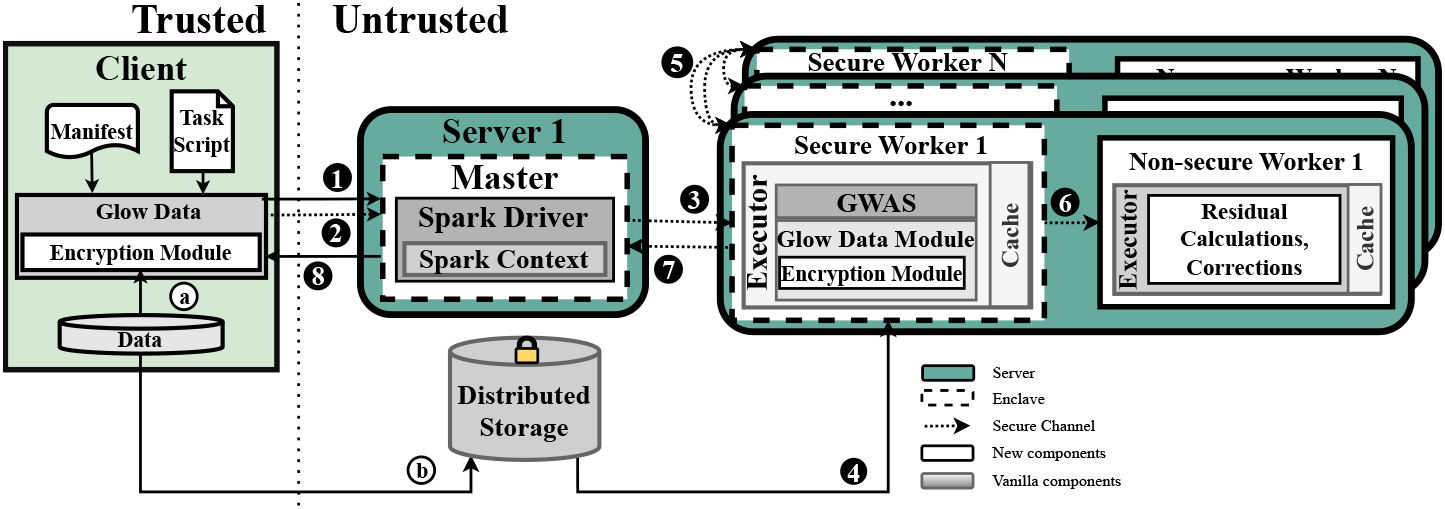
A schematic representation of Gyosa architecture. A Biocenter encrypts and sends its data to distributed data storage. This data is leveraged to perform GWASes. These studies run in a secure and distributed manner with Gyosa.

### Challenges

Users trust external entities to keep their information secure when offloading data storage and processing to third-party infrastructures. However, there have been several reports detailing both external (*e*.*g*., done remotely by a hacker) and internal attacks (*e*.*g*., led by a malicious system administrator with physical access to the cluster) that have successfully compromised the privacy of sensitive information kept at remote infrastructures, such as the one provided by Cloud Computing services [16]. Indeed, these attacks are one of the major barriers limiting the broader adoption of cloud services for storing and processing clinical data, such as genomic data, since its disclosure can lead to the complete identification of individuals [21, 28]. For instance, early studies showed that only 75 SNPs could help identify an individual [21]. In addition, while carrying information on disease predisposition, the leakage of genomic data may imply privacy risks for the individual and their future and previous generations [15].

When outsourcing a genomic data processing pipeline (*i*.*e*., data collection, processing, and sharing) to third-party infrastructures, users increase the attack surface and become susceptible to novel attacks. Namely, in membership inference attacks, the adversary (attacker) can leverage the knowledge it has on a specific individual (*e*.*g*., any genetic information, disease predisposition) and query the analysis results explicitly. Specifically, Homer’s attack and its variations use the background information of the human genome currently available in the public domain to infer whether an individual’s genetic variants information was used for a specific study [37]. Re-identification attacks aim to reveal the identities of individuals whose data have been anonymized and used in genomic analysis. Recent studies have shown that demographic information can be linked with genealogical databases to leak private genomic data [22]. Data poisoning attacks have the goal of manipulating analysis results. Namely, these attacks can generate false assumptions and associations when applied to GWASes, introducing bias and yielding erroneous discoveries [26]. For a detailed overview of these attacks and the genomic pipeline, see Section 2 of Supplementary Material.

### Related Work

To enable the execution of GWAS at untrusted third-party infrastructures while ensuring the privacy of data, several approaches have been proposed based on standard cryptographic primitives such as homomorphic encryption (HE), multiparty computation (SMPC) and differential privacy (DP) [19]. HE enables arithmetic operations (*e*.*g*., sums and aggregations) over encrypted data, thus protecting sensitive content when stored and processed. Despite its recent advances, it has been shown that for GWAS, this technique still imposes a high penalty on execution time and can only support a limited number of algorithmic operations [17]. SMPC enables secure data storage and computation even across multiple entities that do not trust each other. However, this technique resorts to distributed protocols that require several rounds of network communication, which adds a significant delay to the execution of GWAS [39]. DP provides a less penalizing solution in terms of performance overhead. However, it lacks robustness since adding noise to the data and computation can compromise the accuracy and undermine the results. Furthermore, Differential Privacy (DP) cannot effectively manage high-dimensional data or cope with growing data volumes [12]. Thus, the presented solutions have limitations that hinder their use when applied to algorithms to perform GWASes, such as Linear and Logistic Regression or the *X*^2^ test, which are considered in this work.

Recently, Trusted Execution Environments (TEEs) (*e*.*g*., Intel SGX [24], AMD Trust-Zone [1]) have emerged as alternative solutions to ensure privacy-preserving computation and storage in untrusted infrastructures for genomic data [20, 9]. They provide a secure memory space at each server, where genomic data can be securely processed in plaintext format. Both external and internal attackers, even with high Operating System (OS) privileges, cannot access this protected region and disclose the data being processed. While this is a promising technology for running GWAS securely, its application has been typically limited to a single-server mode [5, 18, 32]. By taking advantage of distributed infrastructures, it is possible to enhance the speed and scalability (*i*.*e*., the amount of data being analyzed) of GWASes. However, as highlighted in this paper, developing a distributed solution for privacy-preserving GWAS requires a fundamentally new design that differs from previous methodologies. Namely, it requires securing both the computation and storage of data at each cluster server and the data being exchanged across servers.

### Our Contributions

We propose Gyosa, a novel distributed and privacy-preserving framework for securely executing GWAS in untrusted distributed infrastructures. Gyosa is built on top of Apache Spark [38], a distributed computation framework, and uses Glow, a library for genomic processing that includes regression-based algorithms, statistical tests, and population stratification methods to perform GWAS easily [13]. These are combined with TEEs, namely Intel SGX, to provide a secure environment where sensitive genomic information can be efficiently processed in plaintext without disclosing it to internal or external attackers. Our approach distinguishes itself from recent proposals such [39, 20], as it promotes the outsourcing of computation in a distributed manner in untrusted third-party entities. Also, by resorting to Glow, Gyosa allows the extension of the current genomic analysis pipeline, in which one can add new tasks (*e*.*g*., new statistical tests, genomic imputation, and querying). Expanding on our previous work [2], our solution enables fine-grained differentiation between sensitive and nonsensitive information processed by GWAS. By securely offloading sensitive information to TEEs, we show that it is possible to run large-scale GWAS-based algorithms while protecting the confidentiality of critical data.

To validate our solution, we ran three algorithmic versions based on Logistic Regression, Linear Regression, and *X*^2^ statistical tests. The first two algorithms were run with a variable workload on the benchmarking dataset *“Genome in a Bottle”* [40]. The *X*^2^ test was run against a synthetic dataset simulating 8 *∗* 10^5^ VCF files (one file per individual) with 1*∗*10^6^ random unique SNPs. The first two tests compare the performance impact of our solution against a baseline setup without any security guarantees. The third experiment is intended to evaluate the scalability, feasibility, and behavior of Gyosa with increasing servers.

First, the results show that Gyosa does not affect the quality of the GWASes results. As expected, there is a trade-off between runtime computation and the level of privacy-preserving guarantees. Gyosa shows a runtime execution over-head ranging from 2.5*X* for *X*^2^ statistic tests to 10*X* on regression-based algorithms. Importantly, the results highlight that it is possible to effectively reduce the computation runtime overhead by 2.7X times when distributing the computation across multiple nodes. Indeed, this is a key takeaway from this work, which shows that by distributing GWAS computations, one can achieve practical and usable privacy-preserving designs. Gyosa’s code is available at https://github.com/claudiavmbrito/Gyosa.

## 2 Methods

### 2.1 GWAS

Genomic pipelines include several tasks, such as Short-read Sequence Alignment, Genome Imputation, Variant Call, and GWAS. This work focuses on the latter, which tests the correlation between an SNP’s allele frequency and the presence or intensity of a specific phenotype. Such studies have been performed for a large number of phenotypes, including many diseases. Tested individuals are separated into case and control groups, and to achieve significant and robust results, large sample sizes are required [36].

#### Statistical Methods

To evaluate the performance of Gyosa, we illustrate its practical application by employing two machine learning (ML) algorithms, Logistic and Linear Regression, and test the association with continuous and binary phenotypes. Following the approach in [20], we used the *X*^2^ test to evaluate the scalability of Gyosa and its performance for different workloads with an increasing number of workers. The choice of such algorithms is corroborated by the current state-of-the-art, which shows that these algorithms are widely used to perform GWASes [36]^4^. Next, we analyze the computational complexity of these algorithms using the Big-*O* notation.

##### Linear Regression

checks the relation between one dependent continuous variable (*e*.*g*., weight or blood pressure) and multiple independent ones. The algorithm is efficiently implemented based on matrix multiplications and inversions. For a matrix of *m* x *n* dimensions, where *m* is the number of samples and *n* is the number of features, the time complexity approximates *O*(*m ∗ n*^2^ + *n*^3^) [10].

##### Logistic Regression

is used to perform binary classification based on dependent variables to distinguish between case and control cases [36]. The time complexity of Logistic Regression can be decomposed into *d* as the size of the phenotype vector and *n* as the covariates or features of the phenotype, *O*(*nd*) [34].

##### X^2^ Test

tests if the observed and the expected frequencies (allele counts) in the case-control groups differ significantly. The test can be defined as follows:

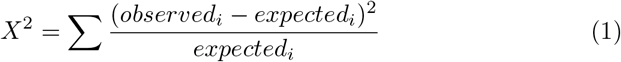

in which we find the observed and the expected frequency of the *ith* SNP. The time complexity of the chi-squared test is *O*(*n*), where *n* is the number of samples [30]. Differently from the previous statistical methods, which intend to test for associations, the *X*^2^ focuses solely on the frequency of the alleles.

### 2.2 Design

Gyosa is a privacy-preserving distributed GWAS-focused solution. It builds on top of SOTERIA [2] and resorts to Apache Spark and Glow for delivering a distributed genomic analysis framework. To achieve the proposed security guaran-tees, Gyosa uses Intel SGX, which provides secure memory regions as enclaves. Apache Spark, Glow, and Intel SGX are further detailed in Section 3 of the Supplementary Material.

#### Threat Model

Gyosa adopts the standard SGX threat model supported by existing research [20, 9, 17]. We consider a scenario where a client seeks to use sensitive genomic data and perform computation on top of it at third-party infrastructures. In this model, the client and the hardware are deemed trustworthy, whereas the third-party infrastructure’s other components (*i*.*e*., host OS, libraries) are regarded as untrusted. This leads to an honest-but-curious adversary model, where the adversary is honest and adheres to the protocol but is also curious and seeks to obtain maximum information.

We highlight that solutions addressing concerns such as *Denial of Service (DoS), side-channel* attacks, or *memory access patterns* can be employed in Gyosa [29]. However, this research is orthogonal to the one proposed here.

#### Gyosa’s Key Components

Gyosa is split into the client module, deployed on a trusted infrastructure, and the cluster module, deployed on an untrusted site (Figure 1). The client module encompasses the encryption of the genomic data and the submission of the GWASes to the untrusted infrastructure. The cluster module includes a distributed Apache Spark and Glow cluster to which the client module will submit the genomic analysis.

##### Client Module

The client module includes three main operations. First, it allows the encryption of VCF files based on authenticated encryption. This mechanism is added to Glow by providing a new class to encrypt this data type. Sensitive data is transparently encrypted before leaving the trusted premises. No changes in the implementation of the analysis scripts are required, thus avoiding any changes to the way users implement or specify their GWAS. Second, it handles the secure outsourcing of the GWAS requests issued by users to the Glow/Spark Cluster. The third operation allows the decryption of the analysis results when returned to users, which are encrypted to avoid revealing sensitive information. To perform a GWAS, users must specify a **task script** file where the analysis steps and all the required parameters are defined. This file contains sensitive information about the analysis task that cannot be leaked or tampered by malicious adversaries. Thus, the encryption module encrypts the task script, which can be sent via unprotected network channels.

The **manifest** is a predefined file that contains the libraries to run the Glow pipeline and the path for the dataset. To ensure data security and integrity when exchanging this file, a secure channel between the client and the Glow/Spark Cluster is created at the analysis bootstrap phase. This secure channel is also used to transmit the user’s encryption key, which is then used by SGX enclaves.

##### Cluster Module

The cluster follows Spark’s workflow on the untrusted site, with a master and *N* workers running on distinct servers. The master is deployed inside an SGX enclave at the untrusted server since the Spark Driver and Spark Context modules require reading plaintext information from the task script to distribute the processing tasks to the workers.

We follow the approach proposed in SOTERIA [2] and deploy a secure and non-secure worker at each of the remaining cluster servers. The secure worker runs inside an SGX enclave and handles all the computation over sensitive data, while the non-secure worker handles non-sensitive data and runs outside of SGX. The exchange of sensitive information between secure workers and the master and between secure workers is done via secure network channels (Figure 1).

##### Partitioned design

Since SOTERIA targets Machine Learning workloads, which significantly differ from GWASes, Gyosa must redefine how computation is partitioned across secure and non-secure workers.

In Gyosa, non-sensitive computations include the residual values (*e*.*g*., matrix calculation of metadata or calculations over single genotype information) and the correction of statistical tests, which in Glow are based on the Firth’s approximation algorithm [23]. In detail, this last step is performed as a score test, which compares the predicted values with the observed values to validate the resulting P-values. All the remaining operations (*e*.*g*., read operations on top of the VCFs, dataframe transformations, and regression operations, among others) are done over sensitive data and performed in SGX enclaves at the secure workers. Note that all this sensitive information remains fully encrypted when transmitted and stored outside of secure workers to ensure its privacy.

In summary, Gyosa differs from SOTERIA by considering a different processing pipeline, namely for GWAses. This implies supporting a new framework (*i*.*e*., Glow), including transparent encryption of a new type of dataset file format (*i*.*e*., VCF files), and redefining the sensitive analysis steps that must be performed inside secure enclaves. Next, we detail the flow of operations in Gyosa.

#### Workflow of Gyosa

In Gyosa, the client ⓐ resorts to Gyosa’s encryption module to encrypt the VCF files at the trusted premises. Then, ⓑ encrypted data is sent to a distributed data storage shared by various servers on the untrusted infrastructure. Similarly, the client specifies the studies it wants to run as task scripts and encrypts these before sending them to the untrusted infrastructure ➊. The default usage of Gyosa assumes a bootstrapping phase between the client and master in which a secure channel is established and used to share the manifest file and the client’s key, which was used to encrypt the VCF files and tasks’ scripts. This key is also used to decrypt the final results ➋.

Following a master–worker architecture, the master, running inside an SGX-enabled server, receives the task the client wants to perform and the path (within the manifest file) for the encrypted dataset. The former is sent encrypted through an insecure channel, while the latter is forwarded inside the previously established secure channel. After decrypting the task script inside the secure enclave, the master forwards specific sub-tasks to each secure worker ➌ through secure channels established between their enclaves. With these sub-tasks, secure workers can fetch the required data from the distributed storage backend and perform the computation. Since data is encrypted at the storage backend, it must be fetched and decrypted at each secure worker enclave to be processed in plaintext (i.e., inside an SGX enclave) ➍.

Following a distributed computation paradigm, workers broadcast intermediate results between them ➎. In addition, following the partitioning of computation, secure workers broadcast metadata to non-secure workers. Again, after performing their computation, the non-secure workers broadcast the information back to the secure workers ➏. Sensitive information shared across secure workers is done through secure channels established between their enclaves. In the final worker-related stage, a consensus regarding the result is reached and sent to the master’s enclave through a secure channel ➐. Final results are aggregated and encrypted by the master, with the client’s key, and sent to the client for transparent decryption at the trusted premises ➑.

## 3 Results

Gyosa was evaluated to understand the impact of adding privacy protection on top of a baseline stack composed by Apache Spark and Glow, which does not provide such guarantees. Two main questions were asked in this evaluation: *a)* How does the execution time of Gyosa compare with the non-secure baseline setup? *b)* How does Gyosa behave when increasing the size of the workload and the number of servers?

### 3.1 Testbed

#### Dataset

For the benchmark, we used a real-world dataset by the *Genome in a Bottle* Consortium [40], with genomes sequenced as part of the Human Genome Project, namely, data from the Ashkenazim Trio family (father, mother, and son). Phenotype information was simulated with the PhenotypeSimulator [25]. The algorithms were tested for different workloads by scaling the original dataset by several factors to reach sizes 1, 4, 16, and 32 GB. For the scalability tests, we generated a synthetic dataset of 80,000 VCF files with 1 *∗* 10^6^ unique SNPs, each VCF file representing one individual. We define the workload size as 20k, 40k, 60k, and 80k individuals.

#### Experimental Setup

Tests were performed in a cluster of 4 servers with OS Ubuntu 18.04.4 LTS and Linux kernel 4.15.0. Each machine has a 10Gbps Ethernet card connected to a dedicated local network and 16 GB of memory. Gyosa uses Apache Spark 3.2.1 and was deployed with version 2.6 of the Intel SGX Linux SDK with driver 1.8, with 4 GB of memory. The client and Spark master run on one server, while Spark workers are deployed on the remaining servers.

### 3.2 Secure GWAS

To assess the impact of Gyosa’s security mechanisms on the execution time of the algorithms, we compare the results with the ones for the baseline setup.

Figure 2 shows the Linear Regression and Logistic Regression algorithm results. In detail, in Figure 2a, for a workload of 1 GB, the runtime overhead of the linear regression algorithm is around 4.5x. For 32 GB, the overhead of Gyosa reaches the maximum for the performed tests, with 10x compared to the baseline setup. The comparison of probability values (*p-values*) between the two approaches shows negligible differences (see Figure 2b). The slightly different values observed may result from the final approximation since it deals with small values and reverts them to *−log*_10_(*p − value*). Similarly, for the Logistic Regression algorithm and workload size of 1 GB and 32 GB, Gyosa shows a runtime overhead of 4x and 9.5x (see Figure 2c).

**Fig. 2:**
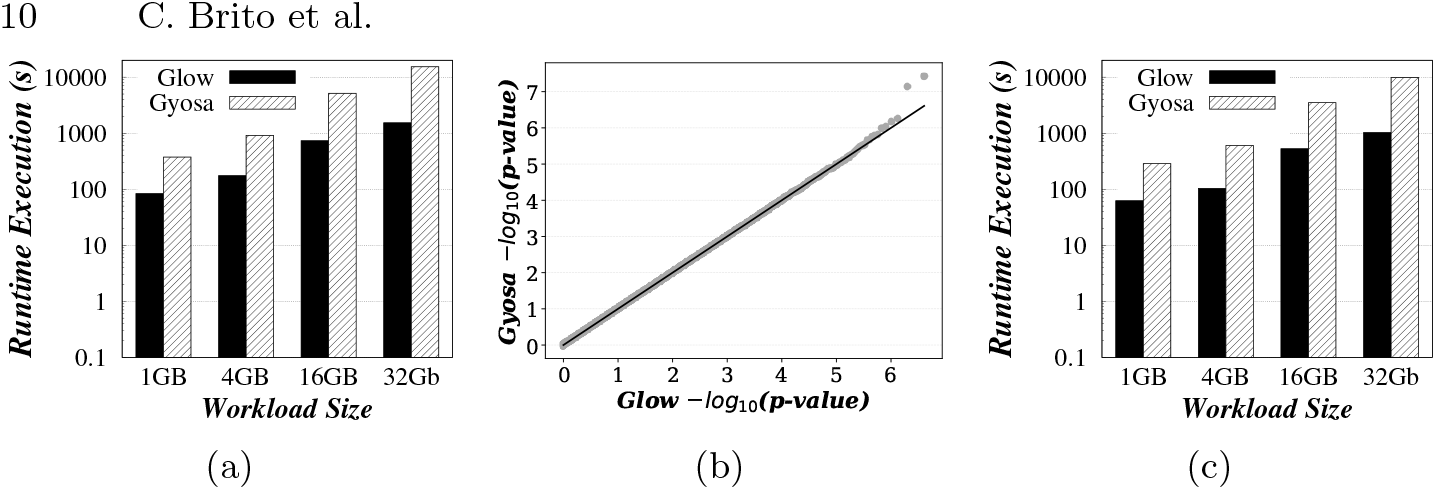
The impact of Gyosa on variable workloads (1GB, 4GB, 16GB, and 32GB) with the Linear Regression and Logistic Regression algorithm compared to baseline. *a)* Execution time of Linear Regression in logarithmic scale; *b)* Comparison of p-values between approaches for Linear Regression; *c)* Execution time of Logistic Regression in logarithmic scale.

Gyosa can be used in a cluster setup with multiple servers, each including one secure worker and one non-secure worker. Figure 3a shows the results of the overhead imposed for the *X*^2^ frequency test. Gyosa leverages the scalability offered by Apache Spark and Glow, as shown by the experiments with up to 3 servers. We verify a linear decrease in the runtime execution when increasing the number of servers for both Gyosa and baseline setups. Namely, when comparing the execution time obtained by running this experiment over 80,000 VCF files with one and three servers, the runtime execution decreases up to 2.7X or up to 2.4 hours. Regarding the security guarantees, the runtime overhead ranges from 1.3x to 3x for a workload of 40k individuals with three servers.

**Fig. 3:**
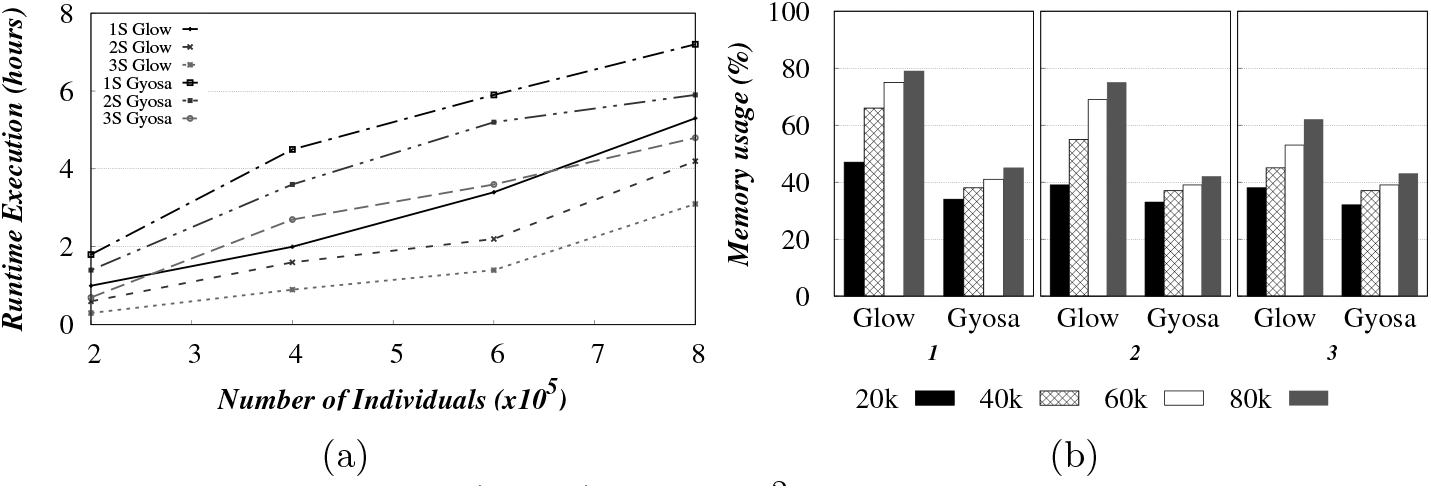
Runtime execution (hours) of the *X*^2^ statistic test for 1 up to 3 servers, supporting workers, and 20k to 80k VCFs from the synthetic dataset. a) Runtime Execution for *X*^2^ statistical test in hours; b) Memory Usage in percentage (%) for *X*^2^ statistical test.

## 4 Discussion

*Security Analysis*. Gyosa combines different mechanisms to safeguard users from attacks (see Section 2 of Supplementary Material for more details on these attacks). It provides transparent authenticated encryption, which protects sensitive data from being disclosed to unwanted parties and ensures anti-tampering properties for clients’ data stored in untrusted infrastructures. This feature protects users from poisoning attacks by limiting access to the plaintext data and not allowing the addition of poisoned data. Membership inference and reidentification attacks are subject to an attacker’s previous knowledge of the genomic data. By ensuring that private data, while at rest and in transit, is always encrypted and that any sensitive computation is performed on SGX enclaves, Gyosa avoids disclosing such knowledge to attackers.

Partitioning the computation across the secure and non-secure workers improves the performance but increases the attack surface. However, previous work shows that genomic data cannot be inferred from the information leaked from sharing metadata and statistical information [27]. Given this assumption, and by not changing the main security protocol specified by SOTERIA, Gyosa can keep the information leakage contained to avoid the success of the aforementioned attacks. The full proofs for SOTERIA’s protocol, as followed by Gyosa, are available at https://dbr-haslab.github.io/repository/sac23.pdf [2].

## Performance and Scalability Analysis

By increasing the number of servers, one can distribute and parallelize GWAS computations and substantially reduce the runtime and memory usage, as observed for the *X*^2^ test in Figure 3a and 3b. Figure 3b shows that despite Glow being a memory-intensive solution [13], increasing the number of servers leads to a decrease in the mean memory usage.

Compared with other SGX-based solutions, namely [4], Gyosa shows comparable runtime overhead. Notably, Gyosa distinguishes from all these state-of-the-art solutions [20, 9] by allowing distributed computation across several servers in a cloud environment.

High-end servers used by state-of-the-art solutions (*e*.*g*., a configuration with *>* 40 cores, *>* 2.0TB of physical memory, and 10TB of disk space [9]) are not widely available and require substantial resources for their setup and maintenance. A cost-efficient alternative is to use cloud environments that allow the distribution of computation by relying on several servers, reducing the execution time of genomic analysis. In Google Cloud [6] with 391.35€ per month, one could opt for *i)* one server with 16 cores and 64 GiB of memory or *ii)* four servers with four cores each and 16 GB of memory. While solution *i)* provides 730 h of computation, solution *ii)* provides a total of 2, 920 monthly hours of computation [14].

Currently, the second generation of SGX includes a page cache with 128 MB. Data that does not fit on this cache must be swapped to/from an encrypted memory region, leading to a performance penalty [11, 8]. Also, in our setup, the amount of encrypted memory attributed to SGX is limited to 4GB in each server. For memory-intensive GWAS algorithms, this means additional swapping to/from disk [24, 11]. Note that this SGX technological limitation cannot be solved by simply upgrading the server’s hardware but is addressed by Gyosa’s distributed design. Namely, by distributing computation across multiple servers, our solution can leverage the aggregated page cache and encrypted memory sizes of multiple servers, which justifies the decreased runtime execution observed for Gyosa in Figure 3a.

## 5 Conclusion

Gyosa offers the first end-to-end privacy-preserving genomic data analytics solution built on top of Apache Spark and Glow. Distributing GWAS computation across multiple untrusted servers allows researchers to efficiently study larger amounts of sensitive genomic data.

Further, by following a computation partitioning scheme tailored for GWAS, Gyosa decreases the amount of data transferred and processed at secure enclaves, which allows for boosting the algorithms’ performance while not compromising security or affecting the quality of the analysis outcomes.

Finally, Gyosa stands out from other solutions by enabling the addition of new tasks (*e*.*g*., statistical tests, genomic imputation, and querying) in the genomic pipeline, making it easier to extend the secure analysis pipeline.

## Footnotes

4 We acknowledge the usage of Principal Component Analysis (PCA), commonly deployed for population stratification. Although addressed in [2] for a different use case, PCA is not currently implemented in Glow, and its privacy requirements may differ from the Gyosa current implementation due to its increasing data exchange and convergence rate.

