## Supplemental Material for "A Distributed Computing Solution for Privacy-Preserving Genome-Wide Association Studies"

### GYOSA Supplementary Material

#### 1 Use Case - Deployment Setting

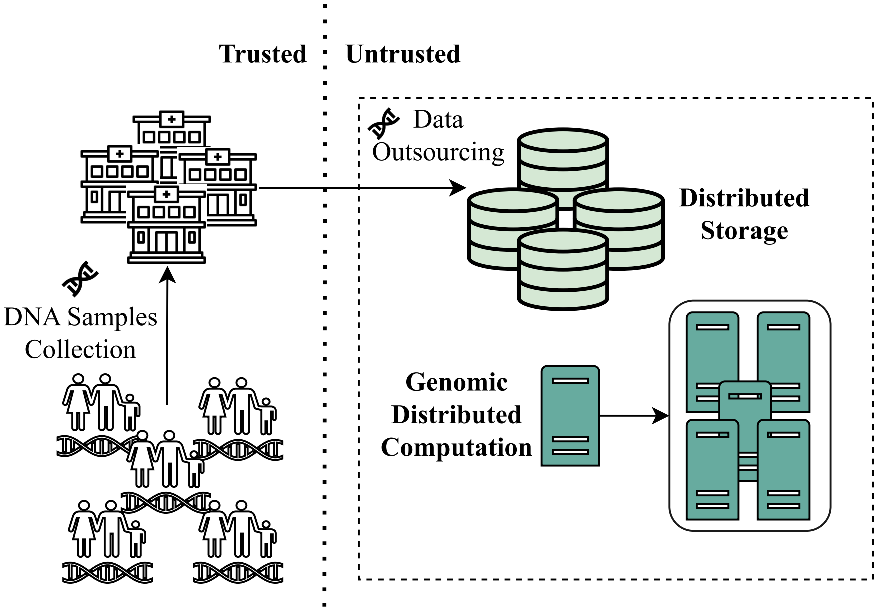


Figure 1: An example where a consortium of entities (*e.g.*, a group of Research Hospitals) have access to a large genomic dataset but lack the processing and storage power to analyze it. In this scenario, the entities can offload the processing and storage of the data to a third-party infrastructure (*e.g.*, a Cloud Computing service) and execute the genomic computation remotely.

#### 2 Background

##### 2.1 Genome

The genome refers to the entire genetic material of a living organism, contained within DNA and shared across all cells. In humans, the DNA sequence is composed of nearly $3.1\text{x}10^9$ base pairs. On average, between each human, the genome presents a variation of $4\text{x}10^6$ base pairs. These differences are commonly called Single Nucleotide Polymorphism, or SNP for short. Based on the reference and alternative alleles from the two copies of each chromosome, SNPs are often depicted in three configurations based on the alleles: 1) aa: reference-homozygous, 2) aA: heterozygous, and 3) AA: alternative-homozygous. SNPs can be detected with techniques like SNP arrays or genome sequencing [2].

##### 2.2 The Genomic Pipeline

Genomic data processing follows a typical pipeline (see Figure 2) that can be divided into three main stages: *i)* data collection, *ii)* data analysis, and *iii)* data sharing.

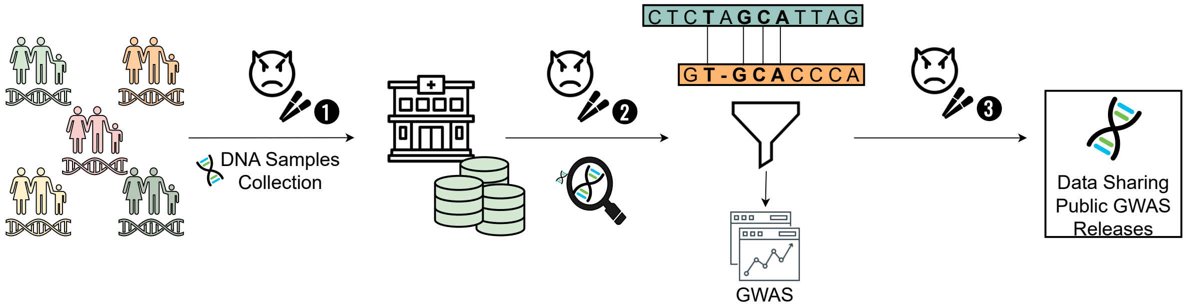


Figure 2: A genomic processing pipeline representing the several steps of the genomic analysis. From the collection of the genomic data to its storage, analysis, and public release. It also represents the several stages where data is most vulnerable ❶-❸.

**Data Collection.** In this stage, the tissue samples are collected from the patient and stored in a biobank. The samples are then processed and analyzed to obtain the genomic data that will be stored in the Biocenter[1] or at third-party (*e.g.*, cloud computing infrastructures) premises according to the imposed regulations over such data ❶.

**Data Analysis.** Data is analyzed to derive useful information. This stage is particularly vulnerable to attacks since the data is usually analyzed in plaintext, which means data is fully open to malicious users while being processed ❷.

**Data Sharing.** In this stage, the data is shared with the patient or the healthcare provider. This stage is vulnerable since data is often shared in an open format and already provides the association between the patient identity and the genetic information ❸.

The existence of sensitive data in a plaintext format at different steps of the pipeline makes it particularly vulnerable to different types of attacks. We now define these vulnerabilities and address the most common attacks that can be performed in the different stages.

##### 2.2.1 Attacks on the Genomic Pipeline

In the deployment of a pipeline on a distributed environment, such as the one in Figure 2, one can expect the attack surface to increase since sensitive data storage and processing are now deployed across multiple servers. However, the key attacks possible in a distributed pipeline are similar to those in a centralized pipeline, where data and computation are outsourced to third-party infrastructures. The three most common attacks are *Membership Inference*, *Re-identification*, and *Data Poisoning*.

1. **Membership Inference** attacks intend to disclose the usage of a specific individual's data in the analysis. The attack is usually performed by an adversary with access to the data and the results. The adversary then tries to infer which individuals were used in the analysis. This is usually done by querying the analysis results for a specific individual and comparing those from other individuals. If the results differ, the adversary can infer that the individual was not used in the analysis. Specifically, Homer's attack and its variations leverage the background information of the human genome that is currently available in the public domain to query models regarding a specific individual [7, 13].

2. **Re-identification** attacks try to reveal the identities of the individuals whose data has been anonymized and used in the analysis. Recent studies have shown that demographic information could be linked to public genomic data and databases. Similarly, it was shown that this could be achieved by linking the genomic data with genealogical databases [8].

[1] We define a Biocenter as any entity that gathers/collects genomic data (e.g., Hospitals, Medical Research Centers).

3. **Data Poisoning** occurs when malicious users want to poison the data in order to produce false results or contaminate the final analysis, usually by adding false data. Data poisoning in the context of genomic data and, more specifically, GWAS, can lead to false assumptions and associations that can create bias and release false insights [10].

### 3 Key Technologies

#### 3.1 Apache Spark and Glow.

Apache Spark is a distributed framework that redefines the programming model of MapReduce to run it in memory [15]. It can be deployed in a cluster of servers and provides a hierarchical architecture composed of a master and several workers; see Figure 1 in the main article. The master node presents two main components: the Spark Context and the Spark driver. Specifically, in this architecture: *i)* the master node receives the tasks (*e.g.*, GWASes, statistical tests) from the client and creates a Spark Context, which then broadcasts the tasks to the workers; *ii)* the workers perform the defined computations and share intermediate results among them; *iii)* when the workers finish their computation, the results are sent back to the master, which aggregates all the results and sends them to the client. The workers to which the master sends tasks are defined by a resource manager who evaluates the available resources and chooses the best possible cluster setting for a specific task [15]. Spark provides a native Machine Learning library (MLlib), a graph library (GraphX), a streaming library (Spark Streaming), and a library for SQL queries (SQL and DataFrames) [12]. The modularity of Spark allows easy integration with several third-party applications. Solutions such as TensorflowOnSpark [14], Glow [6], or ADAM [1] have been built on top of Spark.

Glow offers a solution for performing GWAS at scale. It supports different data formats, including VCF, BGEN, and Plink, by internally converting them into Spark's DataFrames. It includes regression-based algorithms, statistical tests, and population stratification methods to perform GWAS easily [6].

#### 3.2 Intel SGX

Intel SGX provides a set of instructions that offer hardware-based trusted and protected memory regions called *enclaves*. Code and data residing in enclaves are isolated from the host system, increasing the security of their content, which can be handled in plaintext [9, 11]. However, sensitive data and computational outputs transferred to and from the enclave must be encrypted, as these may be handled by untrusted host software. First, the input (ECalls) and output (OCalls) enclave operations often lead to a performance overhead [4, 16]. Then, SGX's enclave has a limited memory capacity (128MB per CPU), which may result in memory swapping that penalizes performance [3]. Even though the second generation of SGX has improved the size of the protected memory region, the Enclave Page Cache (EPC) is still defined as 128MB per CPU [5]. Memory swapping occurs when such a limitation is met, which is a performance-costly mechanism [4].

Therefore, in order to maximize performance, solutions that strike a balance between I/O operations and the volume of data handled by the enclave are required.

[8] Bradley Malin. Re-identification of familial database records. In *AMIA annual symposium proceedings*, volume 2006, page 524. American Medical Informatics Association, 2006.
